# Pivot-and-bond model explains microtubule bundle formation

**DOI:** 10.1101/157719

**Authors:** Marcel Prelogović, Lora Winters, Ana Milas, Iva M. Tolić, Nenad Pavin

**Author notes:** Corresponding authors: N.P. and I.M.T. These authors contributed equally to this work (Marcel Prelogović, Lora Winters, Ana Milas).

## Abstract

During mitosis, bundles of microtubules form a spindle, but the physical mechanism of bundle formation is still not known. Here we show that random angular movement of microtubules around the spindle pole and forces exerted by passive cross-linking proteins are sufficient for the formation of stable microtubule bundles. We test these predictions by experiments in wild-type and *ase1*Δ fission yeast cells. In conclusion, the angular motion drives the alignment of microtubules, which in turn allows the cross-linking proteins to connect the microtubules into a stable bundle.

## Introduction

During mitosis the cell forms a spindle, a complex self-organized molecular machine composed of bundles of microtubules (MTs), which segregates the chromosomes into two daughter cells (1-3). MTs are thin stiff filaments that typically extend in random directions from two spindle poles (4). MTs that extend from the same pole can form parallel bundles, whereas MTs originating from opposite spindle poles form anti-parallel bundles (5-7). Stability of MT bundles is ensured by cross-linking proteins, which bind along the MT lattice, connecting neighboring MTs. Cross-linking occurs only if the distance between the MTs is comparable with the size of a cross-linking protein. These proteins can be divided into two classes: (i) proteins that cross-link MTs without directed movement along the MT, such as Ase1/PRC1 (ref. (8)); (ii) motor proteins that walk along the MT either towards the plus end of the MT, such as Cut7/Eg5 (ref. (9,10)), or towards the minus end, such as Ncd (ref. (11,12)).

Spindle self-organization was studied in different biological systems and several theoretical models were proposed. Formation of antiparallel bundles of MTs in somatic cells of higher eukaryotes was investigated by computer simulations, which include MTs that grow in random directions from two spindle poles and motor proteins that link them (13). Further, several studies have explored the forces generated in the antiparallel overlaps in vitro (14-16) and in *Drosophila* embryo (17-19). Spindle formation was studied in *Xenopus* eggs, using the “slide and cluster” (20,21) and liquid crystal models (22,23). In budding yeast, it is suggested that MTs growing in arbitrary directions from the opposite spindle poles can change their direction due to minus end directed kinesin-14 motors bound to both MTs and get aligned, forming anti-parallel bundles (24). During spindle positioning, myosin motors walking along actin cables accelerate pivoting of astral MTs when they search for cortical anchor sites (25). Studies in fission yeast have shown that passive (thermal) pivoting motion of MTs around the spindle pole body accelerates kinetochore capture (26-28), together with dynamic instability of MTs (29). MT rotational motion about a pivot at the SPB was also included in the model for spindle formation (30) and in vitro studies (31). However, observation of the dynamics of bundle formation in vivo and a corresponding physical model are required to understand the formation and stability of MT bundles.

In this paper, we combine experiments and theory to explore the formation of parallel MT bundles. We introduce the pivot-and-bond model for the formation of parallel MT bundles, which includes random angular motion of MTs around the spindle pole (26), along with the attractive forces exerted by cross-linking proteins. The model predicts faster bundle formation if MTs diffuse faster and the density of cross-linking proteins is higher, which we tested experimentally. We conclude that the angular motion drives the alignment of MTs, which in turn allows the cross-linking proteins to connect the MTs into a stable bundle.

## Experimentally observed bundle formation

The process of MT bundle formation can be observed experimentally in the fission yeast *Schizosaccharomyces pombe* because of a small number of MTs in the spindle. At the onset of mitosis, two spindle pole bodies nucleate MTs that form the spindle and additional MTs grow from the spindle pole bodies performing angular motion (26). In our experiments, we observed that MTs growing at an angle with respect to the spindle eventually join the bundle of spindle MTs (Supporting Fig. 1a, Supporting Movie 1). Such events are also accompanied by an increase in the tubulin-GFP signal intensity in the spindle, suggesting an increase in the number of MTs in the spindle and arguing against the scenario in which one of the MTs depolymerized (Fig. 1b). Additionally, we used cells with GFP-labeled Mal3, a protein that binds to the growing end of the MT (32). Here we observed MT bundling at a finer time resolution (Supporting Fig. 1c in Supporting Note) and the increased Mal3 signal in the spindle after bundling (Fig. 1d). Aside from MTs joining the already formed spindle, we also observed bundling between pairs of MTs which were both freely pivoting (see Supporting Fig. 1a). We did not observe un-bundling events after the bundles were formed. In all cases, MTs performed angular motion around the spindle pole body, which allowed them to approach each other and form a bundle.

**Fig. 1.**
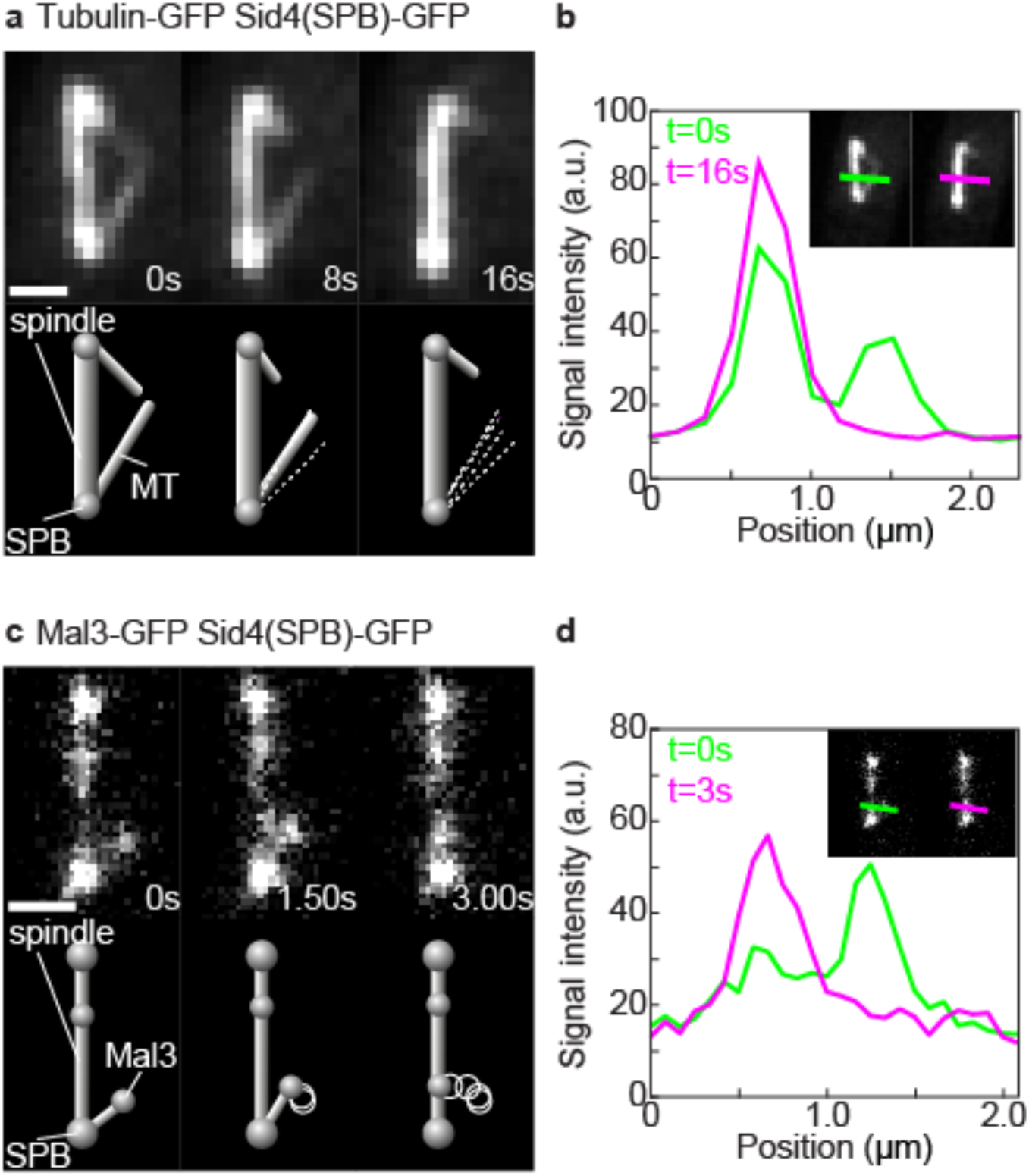
Formation of MT bundles in *S. pombe* cells. (a) Time-lapse images and the corresponding drawings showing the formation of a parallel MT bundle in an *S. pombe* cell expressing tubulin-GFP and Sid4-GFP. (b), Measurement of the tubulin-GFP signal intensity of MTs before bundling and after bundling (measurement done along the line in the inset with the corresponding color). The measurements were done on the first and the last image in panel a, respectively. (c), Time-lapse images and the corresponding drawings showing the formation of a parallel MT bundle in an *S. pombe* cell expressing Mal3-GFP and Sid4-GFP. (d), Measurements of the Mal3-GFP signal intensity of the spindle and MT before bundling as in b. Scale bars in panels a and c are 1μm.

## Theory

To explore the physical principles underlying the formation and stability of MT bundles, we introduce the pivot-and-bond model (Fig. 2a). We describe two MTs as thin rigid rods of fixed length with one end freely joint at the spindle pole body, based on experimental observations both in vivo (26,27) and in vitro (33). The orientation of the first MT at time *t* is described by a unit vector 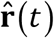 (Fig. 2b). The orientation of the unit vector changes as

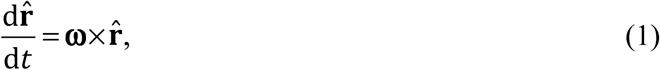

where the vector **ω** denotes angular velocity of the MT. The other MT has a fixed orientation along the z-axis in the direction of unit vector 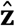. In the overdamped limit, the angular friction is balanced by the torque, **T**, experienced by the MT:

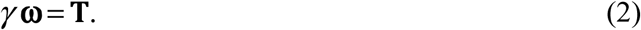

**Fig. 2.**
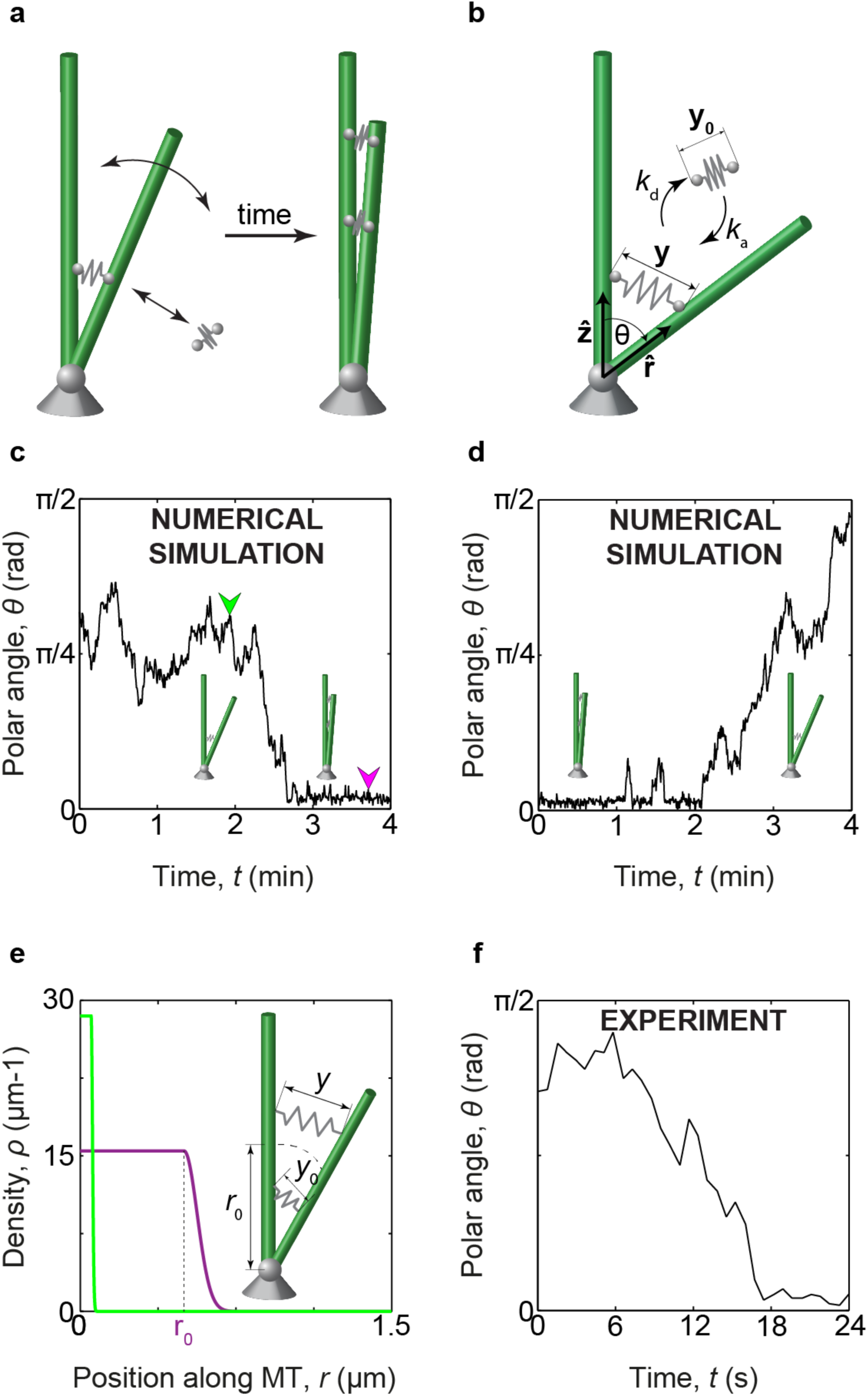
Scheme of the model and numerical solutions. (a), Cartoon representation of the bundling process. A MT (green rod) pivots around the spindle pole body (grey ball). Cross-linking proteins (grey springs) attach and cause the MTs to from a bundle. (b), Scheme of the model. The orientations of two MTs are represented by the unit vectors 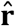 and 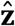. Cross-linking proteins attach to and detach from MTs at rates *k*_a_ and *k*_d_, respectively. The elongation of the attached cross-linking protein is denoted y and their relaxed length is y_0_. The angle between the MTs is denoted *θ*. (c), A sample path for the starting angle *θ* = 0.9 rad, which shows a bundling event. (d), A sample path for an unbundling event. (e), Cross-linking protein density profiles along the MT for two points on the path shown in c, denoted by arrowheads in corresponding colors (large image). The point *r*_0_is shown for the magenta line. The inset shows a schematic of the orientations of attached cross-linkers. The cross-linkers attached within the distance *r*_0_from the spindle pole body are always relaxed, while those attached at larger distances from the spindle pole body are elongated. (f), A sample of MT angle time series obtained using light microscopy on cells with *c* = 300 μm^−1^, *R* = 1.5 μm, *D* = 0.001 rad^2^s^−1^, other the Mal3-GFP label. All calculations are done with parameters shown in Table 1.

**Table 1.** Values of the constant parameters used in this paper.

| Parameter | Value | Source |
| --- | --- | --- |
| $k$ | $0.1 \text{ pNnm}^{-1}$ | Value for Eg5 (ref. (16,39)) |
| $f_c$ | $3 \text{ pN}$ | Value for kinesin-1 (ref. (40)) |
| $y_0$ | $40 \text{ nm}$ | Value for Prc1 (ref. (41,42)) |
| $D_m$ | $0.05 \mu\text{m}^2\text{s}^{-1}$ | Value for Ase1 (ref. (15,43)) |
| $k_a$ | $0.01 \text{ s}^{-1}$ | Value for Ase1(ref. (15)) |
| $k_{d0}$ | $0.1 \text{ s}^{-1}$ | Value for Ase1 (ref. (15)) |

Here, *γ* denotes the angular drag coefficient. We calculate the total torque as 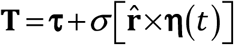, where the first and the second term represent the deterministic and the stochastic components, respectively. If the noise is caused by thermal fluctuations, as in fission yeast (26), **η** = (*η*_*i*_), *i* = 1,2,3 is a 3-dimensional Gaussian white noise, where i-th and j-th components for times *t* and *t*’ obey 〈*η*_*i*_ (*t*),*η*_*j*_ (*t*’) 〉 = δ(*t* –*t* ‘)δ_*ij*_, with δ(*t* – *t*’) being the Dirac delta function and δ_*ij*_ is the Kronecker delta function. The magnitude of the noise is related to the angular drag coefficient, following the equipartition theorem, as 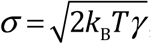, with *k*_B_*T* being the Boltzmann constant multiplied by the temperature. We introduce the angular diffusion coefficient, *D* = *k*_B_*T/γ*, and the equation (2) now reads

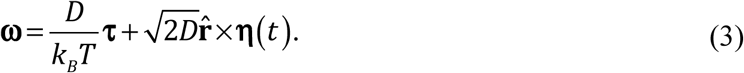

In our model, the torque **τ** in equation (3) is the consequence of forces exerted by cross-linking proteins connecting both MTs. If we denote the positions along the MTs as 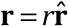 and 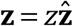 respectively, the torque contribution from cross-linking proteins is

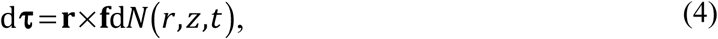

with d*N* being the number of cross-linking proteins connecting the MT elements [*z*, *z* + d*z*] and [*r*,*r* +d*r*]. The force exerted by a single cross-linking protein is elastic and calculated as **f** = –*k* (**y** – *y*_0_ŷ). Here, *k* is the Hookean spring constant, **y** = **z** – **r** is the elongation of the protein linking positions **r** and **z**, with magnitude *y* and direction ŷ= **y/***y*, and *y*_0_ is the relaxation length of the cross-linking protein. We describe the distribution of cross-linking proteins along the MTs by introducing the density, *ρ*, which obeys d*N*(*r*, *z*,*t*) = *ρ* (r, *z*,*t*)d*r* d*z*. To calculate the total torque we summed up all the attached cross-linking proteins:

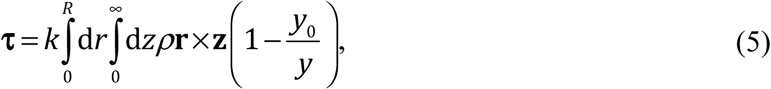

where we used **r** ×(**z** – **r**) = **r** × **z** and allowed the fixed MT to span the entire positive z-axis. When the total number of cross-linking proteins is large we can use the mean field limit and consider them continually distributed along the MT. In this limit, the cross-linking protein density is given by:

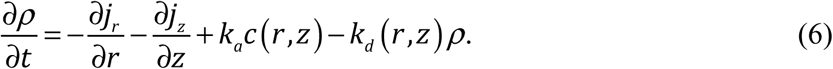

Here, the currents describe the redistribution of cross-linking proteins along the MTs, *j*_r,z_ = *v*_r,z_ *ρ* – *D*_m_∂_r,z_ *ρ*, where the two terms correspond to the drift and the diffusion of cross-linking proteins, respectively. For passive cross-linkers, the velocities are calculated from the balance of the elastic force and friction of cross-linking proteins moving along the MT, *v*_r,z_ = *f*_r,z_/*γ* _m_, where the components of the elastic force parallel with the corresponding MTs are calculated as 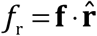 and 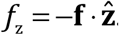. The friction coefficient is calculated using the Einstein relationship, *γ* _m_ = *k*_B_*T/D*_m_. We assume that the attachment rate *k*_a_is constant and that the detachment rate depends on the force experienced by the cross-linking proteins (34), *k*_d_ (*r*, *z*) = *k*_d0_ exp [*f* (*y* (*r*, *z*))/*f*_c_] with *f*_c_ being the critical force required for rupturing the MT-protein bond. The extensions of cross-linking proteins in the nucleoplasm are in thermodynamic equilibrium, hence they obey the Boltzmann distribution. Thus, the distribution of cross-linking proteins in the nucleoplasm with respect to their extensions is given by 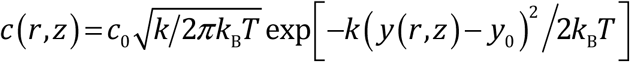, where the constant *c*_0_ is the linear concentration of cross-linkers in the nucleoplasm. Equations (1)-(6) provide a complete description of angular movement for the MT in the presence of cross-linking proteins.

## Results

To obtain the time course of the MT orientation, we parameterize the orientation of the MT given by the unit vector by 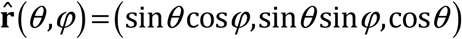, where *θ* and φ denote the polar and azimuthal angle, respectively. In this parameterization, the equation of motion for the polar angle reads

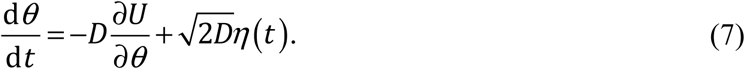

The normalized potential describing the interaction between the MTs, *U* (*θ*), is implicitly defined as –∂_*θ*_*U* =*τ/k*_B_*T* + cot *θ*, where *τ* denotes the magnitude of the torque and cot *θ* is the spurious drift term (35) (for derivation see Supporting Note 1). This equation is sufficient to describe the bundling process, because the angle between the MTs is given only by the polar angle, the dynamics of which is independent of the azimuthal angle. In the adiabatic approximation, ∂*ρ/*∂*t* = 0, equation (6) yields a one-dimensional cross-linker density profile *ρ*_r_ (*r*) (exact expression given by equation S21 in Supporting Note 1). Integrating the density profile over the entire length of the MT allows us to calculate the torque exerted by cross-linking proteins, which in turn allows us to calculate the generalized potential *U* (*θ*) = –*θ*_max_[Θ (*θ* –*θ*_min_/*θ* + Θ(*θ*_min_ –*θ*) *θ*_min_] – ln [sin(*θ*)]. Here, *θ*_max_ = *k*_a_*c*_0_ *y*_0_/4*k*_d0_ is the local maximum of the potential, *θ*_min_ = *y*_0_/*R* is the local minimum and Θ is the Heaviside step function. The expression for the generalized potential combined with equation (7) formulates the pivot-and-bond model in terms of a one-dimensional Langevin equation.

By numerically solving equation (7) for the polar angle, for a large initial angle, we found that the MT performs random movement and spans a large space (Fig. 2c). However, the movement can become abruptly constrained in the vicinity of angle zero. These small angles correspond to a bundled state. Our numerical solutions also show that, in rare cases, constrained MT movement in the vicinity of angle zero can suddenly switch back to free random movement (Fig. 2d). The constrained movement near angle zero is a consequence of short range attractive forces exerted by the cross-linkers that accumulate in larger densities when MTs are in close proximity (compare green and magenta lines in Fig. 2e). The density is constant up to *r*_0_because in that region, the cross-linkers can always attach in a relaxed configuration, while for *r* > *r*_0_, the cross-linkers will always be under tension and their density will drop off dramatically as *r* increases further (see illustration in Fig. 2e and Supporting Note 1). Our numerically obtained time courses that correspond to the MT bundling are similar to those from experiments (compare Figs. 2c and 2f).

To systematically explore the formation of MT bundles and their stability, we first examine the normalized potential describing the interaction between the MTs. The shape of the normalized potential for different MT lengths and nucleoplasm cross-linker concentrations is shown in Fig. 3a. The normalized potential has a local maximum that introduces an intuitive boundary between two regions, *Θ*_B_ ≡[0,*θ*_max_] and *Θ*_u_ ≡[*θ*_max_, *π*], termed bundled and unbundled state, respectively. The bundling and unbundling probabilities are calculated as 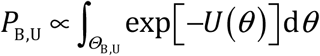, where we used the stationary probability distribution exp[–*U* (*θ*)]and the normalization *P*_B_ + *P*_U_ = 1 (see ref. (35)). Bundling probability, shown in Fig. 3b as a function of cross-linker concentration, exhibits a sharp transition from zero to one around the value *P*_B_ = 1 2. Based on this transition, we define bundles as stable if *P*_B_ > *P*_U_. Thus, our theory predicts that stable bundles can form only if MTs are long enough and there are enough cross-linking proteins in the nucleoplasm (see Fig. 3c).

**Fig. 3.**
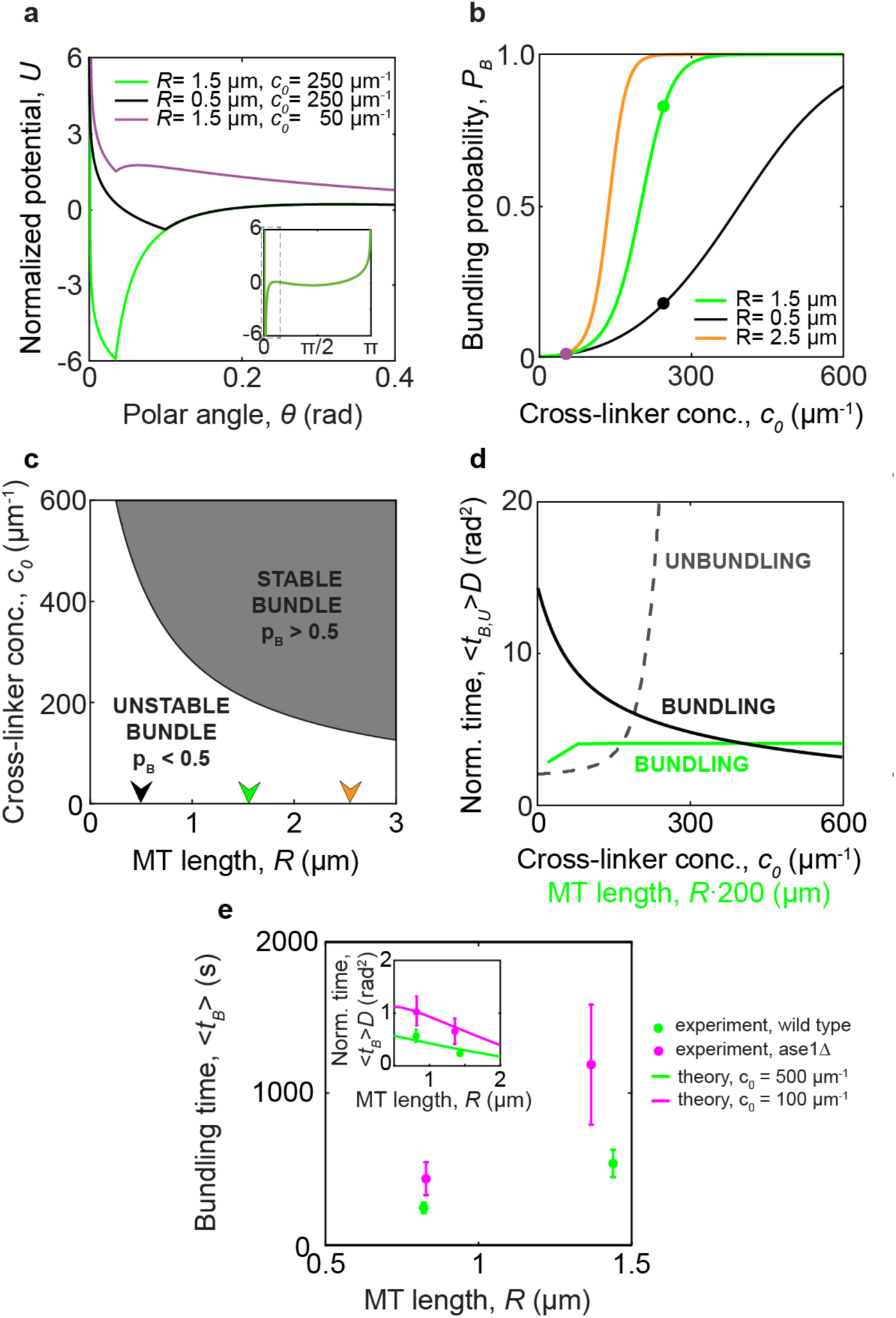
Solutions of the model in the adiabatic approximation. (a), The effective potential as a function of the polar angle for small angles. The inset shows the potential for all angles. (b), Bundling probability as a function of *c*_0_for three different values of *R*. The colors of the dots correspond to the color code of the parameters used in a. (c), Phase diagram. The gray area represents the region where bundles are stable. The arrowheads represent the values of *R* used in b. (d), Normalized bundling and unbundling times. The black lines represent the normalized bundling (solid line) and unbundling time (dashed line) as a function of *c*_0_for *R* = 1.5 μm. The green line represents the normalized bundling time as a function of *R* for *c*_0_ = 250 μm^−1^. (e) Experimentally measured bundling times for wild type (green) and *ase*1Δ cells (magenta). Inset shows the comparison between measured normalized bundling time (dots with error bars) and theoretical curves for *c* = 500 μm^−1^ (solid green line) and*c*_0_ = 100 μm^−1^ (solid magenta line).

Finally, we calculate how the MT bundling time depends on the parameters of the system. In the case of an isotropic distribution of initial MT orientations, we calculate the bundling time as 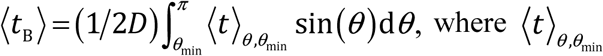 is the first passage time from an initial angle *θ* to the angle *θ*_min_ (for more details see Supporting Note 1). After solving these integrals numerically, we found that the bundling time normalized by the diffusion coefficient, 〈*t*_B_〉 *D*, decreases as the cross-linker concentration increases (solid black line in Fig. 3d), but is not significantly affected by the MT length (solid green line in Fig. 3d). Note that the bundling time, 〈*t*_B_〉, is inversely proportional to *D*, which decreases with MT length (26), thus we expect that the bundling time increases with MT length. The unbundling time, 〈t_U_ 〉, is calculated analogously. The unbundling time becomes longer than bundling time if the condition for bundle stability *P*_B_ > *P*_U_is fulfilled. Once this condition is satisfied, the bundling time greatly increases (dashed line in Fig. 3d).

In order to compare the theory with experimental observations, we measured the bundling time as the total observation time of MTs divided by the number of observed bundling events, 〈*t*_B_〉 = *t*_exp_/*n* (see Supporting Note 2). Along with the wild type cells, we also performed the measurements on the mutant in which the ase1 cross-linker was knocked out (denoted *ase*1Δ, ref. (36,37)), in which we also observed MT bundling (see Supporting Fig. 1b, Supporting Movie 4 and Supporting Table 1). We observed that the bundling time increases with MT length (Fig. 3e), and that the bundling time is significantly longer in *ase*1Δ cells (compare green and magenta line in Fig 3e). We normalized the bundling time by the diffusion constant and found a weak dependence on MT length, but a significant increase in *ase*1Δ cells compared to wild type (inset of Fig 3e, for theory see Supporting Note 1, for diffusion measurements see Supporting Fig. 2). The theory reproduces the weak dependence on MT length and implies that the deletion of ase1 decreases the effective cross-linker concentration roughly five fold.

In conclusion, our work implies that only passive processes, thermally driven motion of the MTs and passive cross-linkers, are sufficient to describe the formation of parallel MT bundles. By introducing the pivot and bond model we gain a deeper understanding of the mesoscopic properties of the bundling process, such as bundle stability and average bundling time, as well as predict their dependence on biological parameters such as MT length and cross-linker concentration. This approach is complementary to more exhaustive and detailed methods such as large-scale numerical simulations (for example (30,38)).

Along with parallel MT bundles, mitotic spindles also contain bundles of anti-parallel MTs, which are made of MTs extending from the opposite spindle poles. The theory developed here could be generalized to describe MTs extending from two spindle poles by adding additional angular variables and including directional movement of cross-linking proteins. Just like here, such model will give insight into the minimal requirements for the formation of anti-parallel MT bundles and therefore shed additional light on the physics of spindle formation.

## Author contributions and acknowledgements

N.P. and I.T. designed and supervised the project. M.P. and N.P. developed the theory. L.W. carried out the experiments and A.M. analyzed the experimental data. N.P., M.P. and I.T wrote the manuscript with input from L.W. and A.M. The data that support the findings of this study are available from the corresponding authors upon reasonable request. Contact N.P. for information on the theory presented and I.M.T with requests related to the experimental measurements.

We thank D. Radić and V. Despoja for comments on the manuscript and M. Glunčić for valuable discussions and advice regarding our theoretical model; F. Elsner from the Electronics Service of MPI-CBG for building the thermoelectric device; J. Millar for yeast strains; Light Microscopy Facility of MPI-CBG for help with microscopy; J. Brugués, T.M. Franzmann, A. Garcia Ulloa and the rest of the Tolić lab for discussion and advice; and I. Šarić for editing the figures. We acknowledge the Unity through Knowledge Fund (UKF), German Research Foundation (DFG), European Research Council (ERC) and the QuantiXLie Center of Excellence for funding.

## Supporting references

Ref. (44-47) are used in the Supporting material.

